# Anion-conducting channelrhodopsins with tuned spectra and modified kinetics engineered for optogenetic manipulation of behavior

**DOI:** 10.1101/156422

**Authors:** Jonas Wietek, Silvia Rodriguez-Rozada, Janine Tutas, Federico Tenedini, Christiane Grimm, Thomas G. Oertner, Peter Soba, Peter Hegemann, J. Simon Wiegert

## Abstract

Genetic engineering of natural light-gated ion channels has proven a powerful way to generate optogenetic tools for a wide variety of applications. In recent years, blue light-activated engineered anion conducting channelrhodopsins (eACRs) have been developed, improved, and were successfully applied *in vivo*. We asked whether the approaches used to create eACRs can be transferred to other well-characterized cation-conducting channelrhodopsins (CCRs) to obtain eACRs with a broad spectrum of biophysical properties. We generated 22 variants using two conversion strategies applied to 11 CCRs and screened them for membrane expression, photocurrents and anion selectivity. We obtained two novel eACRs, Phobos and Aurora, with blue-and red-shifted action spectra and photocurrents similar to existing eACRs. Furthermore, step-function mutations greatly enhanced the cellular operational light sensitivity due to a slowed-down photocycle. These bistable eACRs can be reversibly toggled between open and closed states with brief light pulses of different wavelengths. All new eACRs reliably inhibited action potential firing in pyramidal CA1 neurons. In *Drosophila* larvae, eACRs conveyed robust and specific light-dependent inhibition of locomotion and nociception.

## Introduction

The discovery of natural anion conducting channelrhodopsins (nACRs)^1–3^ and the development of engineered anion-conducting channelrhodopsins (eACRs) by targeted mutagenesis of cation-conducting channelrhodopsins (CCRs)^4–7^ introduced a new class of optogenetic tools^8^. The existing eACRs were derived from either *Chlamydomonas reinhardtii* channelrhodopsin-2 (*Cr*ChR2)^7^ or the channelrhodopsin chimera C1C2^5^ using two complementary strategies. Exchange of a single glutamate for an arginine in the central gate of *Cr*ChR2 was sufficient to invert selectivity from cations to anions. Additional exchange of two glutamate residues in the outer pore and the inner gate completely eliminated residual proton conductance, yielding the highly anion-selective eACR iChloC^6^. In parallel, mutation of several amino acids within C1C2 to render the electrostatic potential of the conducting pore more positive, strongly favored anion conductance^5^. Further improvements led to a second highly anion selective eACR iC++ and the related step function version SwiChR++^4^. These improved versions have been successfully used to silence neurons in mice or rats *in vivo*^4,5,9,13^.

We asked whether the approaches used to create eACRs can be transferred to other known CCRs to obtain eACRs with a broad spectrum of biophysical properties, especially different kinetics and spectral sensitivities. So far, all eACRs show action spectra similar to *Cr*ChR2 with maximal activation in the blue spectral range. ACRs with a red-shifted absorption maximum are desirable for three reasons: First, long-wavelength light penetrates deeper into biological tissue due to lower absorption and scattering^14,15^. This enables silencing of larger volumes at reduced light energies compared to blue-light activated tools^16^. Second, combination with blue-light activated tools becomes possible. Third, many animals are blind to light beyond ~600 nm, while visible light exposure can result in positive or negative phototaxis, particularly in invertebrates^17,18^. Red light activation avoids or reduces direct effects of the light pulse on behavior. On the other hand, eACRs with a blue-shifted action spectrum could be combined with red-shifted sensors and actuators, as their excitation would not interfere with activation of such ACRs.

Adding step-function mutations to spectrally shifted eACRs conveys further benefits: In these mutants, a long-lasting chloride conductance can be activated by a short light pulse and terminated by a second light pulse of longer wavelength^4,19,20^. Due to the slow photocycle, photons are integrated over time, increasing the operational light sensitivity of target cells by orders of magnitude.

Here, we report the successful development of two new eACRs, termed Phobos and Aurora, which express well in neurons and provide sufficient photocurrents for efficient silencing. Compared to existing eACRs, Phobos and Aurora exhibit blue - and red-shifted action spectra, making them potentially suitable for dual-wavelength experiments with spectrally distinct actuators or sensors of the optogenetic toolbox. Adding the step-function mutation C128A to Aurora, Phobos and the previously published iChloC^6^ yielded bi-stable versions which could be toggled between open and closed states with short light pulses.

We characterized the biophysical properties of these new eACRs in HEK cells and verified their silencing ability in organotypic hippocampal slice cultures. Potent eACR variants were further tested in larvae of *Drosophila melanogaster*, an organism where the classical light-activated ion pumps halorhodopsin (NpHR)^21,22^ and archaerhodopsin Arch^23^ show modest effects. In contrast to eNpHR, light-activation of Aurora or the step-function variant of Phobos in nociceptive class IV dendritic arborization (C4da) neurons^24^ acutely abolished nociceptive behavioral responses and reversibly decreased locomotion when expressed in motor neurons.

## Results

### Biophysical characterization in HEK cells

To obtain new eACRs with distinct spectral and kinetic properties, we aimed at converting well-characterized CCRs with known biophysical properties into anion channels. For this, two distinct, previously successful approaches were taken^4,5,7^ According to the first approach^7^, we replaced the central gate glutamate (E90R; *Cr*ChR2) with arginine in various CCRs (Figure 1A, B). The new mutants were expressed in HEK cells and tested for membrane expression and photocurrents. *Ts*ChR^E72R^ (*Tetraselmis striata* ChR)^25^, *Ps*ChR2^E73R^ (*Platymonas subcordiformis* ChR2)^26^, *Vc*ChR1^E85R^ (*Volvox carteri* ChR1)^27^ and Chronos^E107R^ (*Stigeocionium heiveticum* ChR)^25^ did not yield detectable photocurrents (Figure 1B, Figure S2A). Because Chronos exhibits no serine at the homologous position of S63, which is a main constituent of the inner gate in *Cr*ChR2 we speculated that a differently arranged inner gate could be responsible for the missing photocurrents. Therefore, we created the double mutant Chronos^A80S E107R^ to reconstitute a serine residue at the putative inner gate while rendering the central gate anion conductive. But this mutant also remained non-functional (data not shown). The chimeric CCRs C1C2^E129R^ ^28,29^ and C1V1^E129R^ ^30,31^ showed partial Cl^−^-conductivity but current amplitudes were below 10 pA and membrane localization was poor (Figure 1B, Figure S2A). The mutated *Tc*ChR (*Tetraselmis cordiformis* ChR)^25^ displayed photocurrents, but the reversal potential was not shifted upon change of the Cl^−^ gradient (Figure 1B, Figure S2A). In case of the spectrally red shifted CCRs Chrimson (*Chiamydomonas noctigama* ChR1)^25^ and ReaChR^32^, the E90R homologous mutation caused reduction of photocurrents and a strong deceleration of channel closing. However, again no (Chrimson) or only minor (ReaChR) shifts of the reversal potential after changing the Cl^−^ gradient were detected (Figure 1B, Figure S2A). The only CCR that could be converted using this strategy was the highly *Cr*ChR2-related *Co*ChR (Figures 1B, S2A and S10) from *Chioromonas* oogama^4,25^. As shown previously, additional introduction of the iChloC homologous mutations (*Co*ChR^E63Q E70R E81S^) further improved the Cl^−^-selectivity (Figures 1B, S3) but neuronal expression revealed toxic side effects^6^. Because the ChloC conversion strategy (i.e. to replace the homologous glutamate 90 in the inner gate) was not generalizable to other CCR variants, we proceeded with the second approach and transferred mutations and an N-terminal sequence from the eACR iC++^4^ to the same group of CCRs (Figure 1A, B). Most constructs (*Tc*ChR, *Ts*ChR, *Ps*ChR2, Chronos and Chrimson) showed no photocurrents and only weak expression and/or membrane localization (Figure S2B). As with the ChloC strategy, *Cr*ChR2^T159C^ and *Co*ChR could be successfully converted and showed high Cl^−^-selectivity. However, both variants showed no improvements compared to iC++ (Figures 1B, S2B) and were not further investigated. Conversion of the red-shifted CCRs^27^ was also successful, but the converted *Vc*ChR1 and C1V1 showed photocurrents below 10 pA. In contrast, the modified *Vc*ChR1-derivate ReaChR displayed robust current amplitudes upon illumination with green light, which were comparable to iC++ photocurrents, evoked by blue-light (Figures 1D,E, S2B). Thus, both engineering strategies previously used to generate iChloC and iC++ are not generally applicable to convert CCRs into eACRs. Only *Cr*ChR2^T159C^, *Co*ChR and ReaChR were successfully transformed with the iC++-based strategy. ReaChR, due to its red-shifted action spectrum, is a promising new candidate for a green/yellow/red light activated ACR. Since no CCR with blue-shifted absorption (*Tc*ChR, *Ts*ChR and *Ps*ChR2) could be successfully converted, we aimed to shift the absorption of iC++ by introducing the mutations T159G and G163A that are present in all three deep-blue absorbing CCRs^25^ (Figure S1). These mutations induced a blue-shift when the homologous mutations were used in the parental C1C2 construct^33^. The double mutant iC++^T159G G163A^ indeed showed blue-shifted maximal activity (Figure 1C) without altering photocurrent amplitudes and membrane expression compared to iC++ (Figures 1D, 3A-C, S2B).

**Figure 1:**
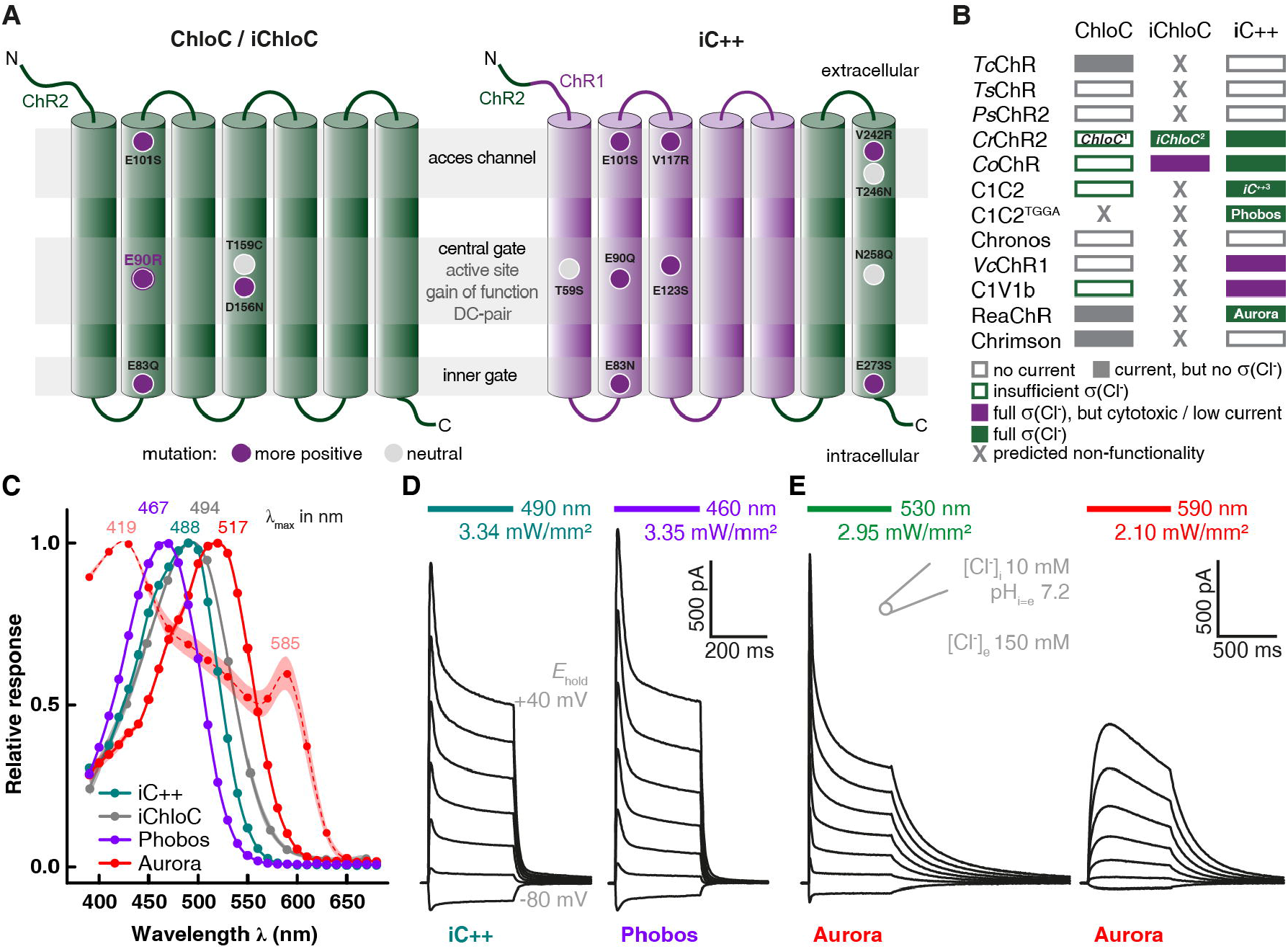
Construct design and screening result for eACRs. **(A)** Conversion strategies yielding the eACRs ChloC, iChloC (left) and iC++ (right). The transmembrane helices of *Cr*ChR2 (ChR2) and CrChR1 (ChR1) are shown in green and purple, respectively. The positions of the outer access channel, central gate (including retinal binding pocket) and inner gate are indicated by gray horizontal stripes. Mutations are displayed as circles at the relative position within the respective helix. ChloC has the mutations E90R and T159C, whereas in iChloC E83Q and E101S were additionally introduced. The D156N mutation from slowChloC is also present in iChloC (left). iC++ exhibits 10 mutations and a modified N-terminal sequence (right). **(B)** Summary of the mutation transfer approach. Most ChR-variants harboring ChloC-or IC++-mutations showed no photocurrents. In addition, of constructs, which produced a photocurrent, the majority had no or only partial Cl^−^ conductivity (ơ(Cl^−^)). For details and ChR abbreviations, please see main text. **(C)** Action spectra of eACRs. Peak wavelengths (indicated above) were obtained from fitting with a 3-parameter Weibull distribution. In addition to typical low intensity action spectra (solid lines) obtained with 10 ms pulsed activation, a spectrum with continuous illumination of 500 ms and tenfold increased photon irradiance was recorded for Aurora (dotted line). Data points show mean ± SEM (n = 6) Phobos, 9 iC++, 7 Aurora, 6 Aurora with tenfold photon irradiance, 6 iChloC). **(D and E)** Typical photocurrent traces at high extracellular [Cl^−^] of the newly developed eACRs Phobos (D, right) and Aurora **(E)** compared to the established eACR iC++ (D, left). The holding potential was increased from −80 mV (bottom) to +40 mV (top trace) in 20 mV steps. Duration of light application at respective wavelengths is indicated by colored bars above the traces.

In summary, our screen of 22 different putatively anion-selective constructs yielded two functional eACRs with novel properties, namely the blue-shifted iC++^T159G G163A^, which we termed Phobos and the red-shifted anion-selective ReaChR variant, which we termed Aurora.

We next characterized the biophysical properties of Phobos and Aurora in HEK cells alongside the two established eACRs iC++ and iChloC. Phobos showed photocurrent properties similar to the parental iC++. Light application evoked fast currents that decayed to a stationary level under constant illumination. After light shutoff the current rapidly decayed to baseline with a time constant of 10.1 ± 0.8 ms (Figure 1D, 2A,F). Aurora, like the parental ReaChR^32^, showed higher inactivation and slower *off*-kinetics (264 ± 19 ms). As previously demonstrated for ReaChR^32,34^, high stationary currents could also be evoked with orange light (590 nm), where no fast peak current is observed, making Aurora suitable for red-shifted activation (Figure 1E).

**Figure 2:**
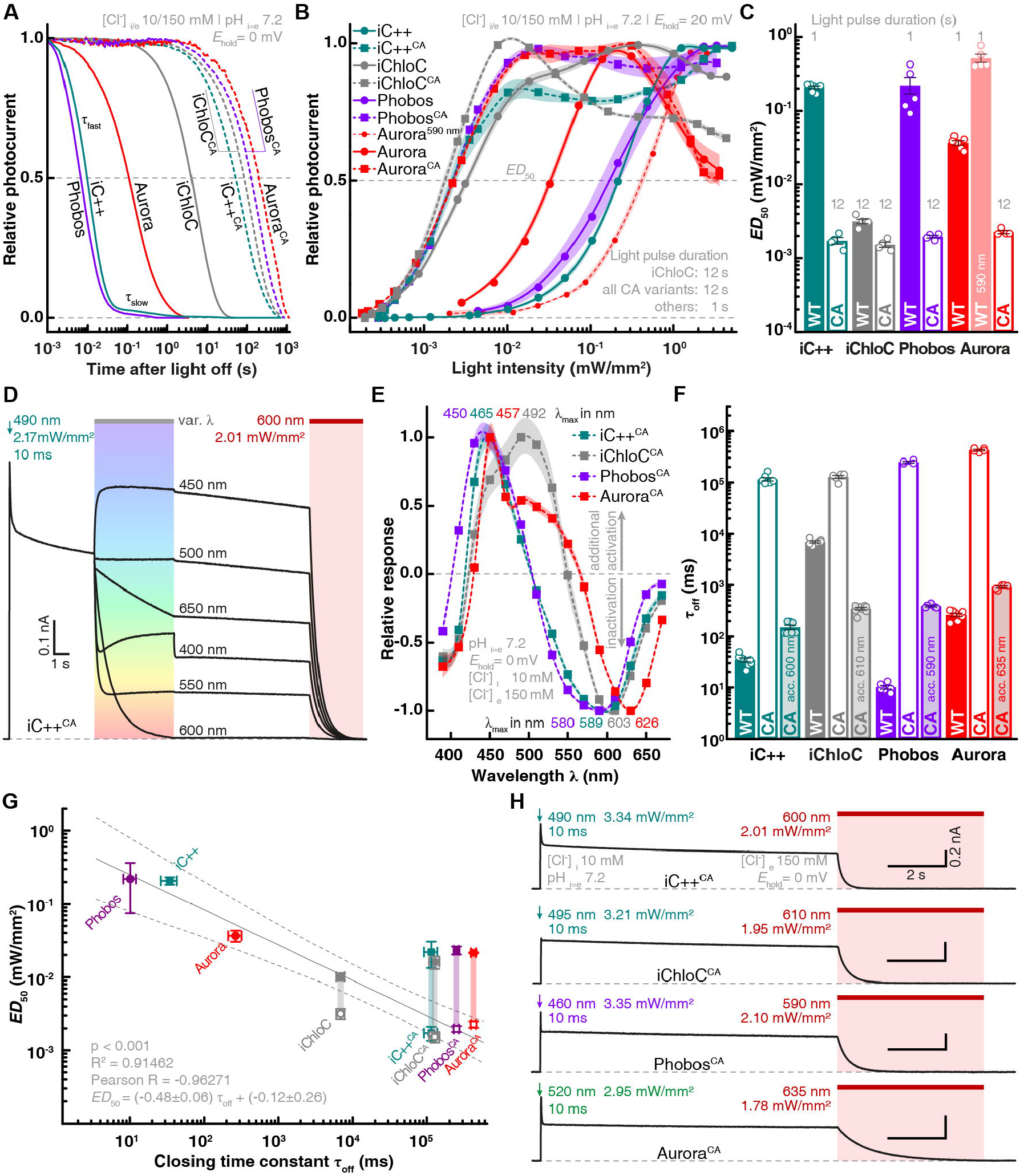
Kinetics, light sensitivity and control of eACRs. **(A)** Normalized photocurrent traces after light shutoff are displayed for fast cycling eACRs and their respective C128A step-function variants logarithmically binned to 50 data points per decade. **(B)** Light titration of eACRs and eACR C128A variants. Stationary photocurrents are normalized to the maximum (n = 5 Phobos, 4 Phobos^CA^, 7 iC++, 6 iC++^CA^, 5 Aurora, 5 Aurora 590 nm, 4 Aurora^CA^, 5 iChloC, 4 iChloC^CA^). 12 s light pulses were used for C128A variants to reach stationary photocurrents. **(C)** Light-activation *ED*_50_ values of fast cycling eACRs (WT) and their C128A variants (CA) obtained from fitted measurements shown in **(B)**. **(D)** Example recording demonstrating the strategy to determine wavelength dependent inactivation for step-function eACRs. After fully opening the channel with a 10 ms light pulse at the peak activating wavelength, light with different wavelengths at identical photon irradiance was applied to accelerate channel closing. The eACR was fully closed with high-intensity red-shifted light after each trial. **(E)** Inactivation spectra calculated from measurements as shown in **(D)**. Positive values show additional activation, whereas negative values denote inactivation (n = 4 Phobos^CA^, 6 iC++^CA^, 5 Aurora^CA^, 5 iChloC^CA^). Wavelengths yielding maximal activation and inactivation are displayed for each eACR C128A above or under the curves, respectively. **(F)** Apparent closing kinetic time constants (ľ_off_) of fast cycling eACRs (WT), their C128A variants (CA) and the accelerated closing by application of red-shifted light. (n = 6 Phobos, 4 Phobos^CA^, 5 Phobos^CA^ accelerated, 8 iC++, 7 iC++^CA^, 7 iC++^CA^ accelerated, 8 Aurora, 5 Aurora^CA^, 5 Aurora^CA^ accelerated, 7 iChloC, 5 iChloC^CA^, 8 iChloC^CA^ accelerated). **(G)** *ED*_50_ vs. closing time constant of eACR variants linearly correlates during extended illumination of C128A variants (12 s) except for iChloC that shows a higher sensitivity compared to other eACRs with respect to its closing time constant. Fitting statistics are shown in the figure panel. **(H)** Typical photocurrent traces of the newly developed step-function eACRs Phobos^CA^, iChloC^CA^ and Aurora^CA^ compared to the established step-function eACR iC++^CA^ (alias SwiChR++), activated by short 10 ms light pulses. Channel closing was always facilitated with red-shifted light. Mean values ± SEM are shown (B, C, E, F and G) together with single measurement data points (dots, C and F).

The action spectrum maxima of the new ACRs were substantially shifted to 467 nm (n = 6, Phobos), and 517 nm (n = 5, Aurora) compared to the maxima of iC++ and iChloC with 488 nm (n = 9) and 494 nm (n = 6), respectively (Figure 1C). Thus, the T159G G163A mutations resulted in a 21 nm blue-shift with respect to the parental iC++. When longer light pulses (500 ms) with 10-fold increased photon irradiance were used, the action spectrum of Aurora broadened^32^, revealing two peaks at 419 nm and 585 nm (n = 6). The central part of the spectrum showed a minimum in the region where the low intensity spectrum was maximal (Figure 1C), possibly caused by inactivation due to absorption of the dark state and the L-state in the same spectral range^34^.

Next, we aimed to slow down the photocycle of iChloC, Phobos and Aurora to create bi-stable eACRs with decelerated closing kinetics^4,19,20^, enabling accumulation of eACRs in the open state during illumination (Figure S4A). As a consequence, under continuous illumination the light dose required to achieve half-maximal photocurrents (*ED*_50_) was orders of magnitude lower compared to non bi-stable eACR variants (Figure 2B, C, G, S4). Thus, the relative light sensitivity (1/*ED*_50_) increased. This gain in ion flux per photocycle allows for optogenetic silencing of cells at low light intensities, if ms-precision is not mandatory^8^. Previously, introduction of the C128A mutation in iC++ produced a bi-stable eACR called SwiChR++ whose closing kinetics can be accelerated by red light, making it switchable between open and closed conformation with short light pulses^4^ (Figure 2D-H). Similar to SwiChR++ (here termed iC++^CA^ for consistency), where we measured a 3333-fold longer closing time constant than for iC++, our new C128A variants of iChloC, Phobos and Aurora showed 19-fold, 24559-fold, and 1604-fold reductions in their *off*-kinetics, respectively (Figure 2A, F). *ED*_50_ values and closing time constants were linearly related at saturating light pulse duration (1 s for Phobos, Aurora and iC++, 12 s for iChloC, iC++^CA^, Phobos^CA^, iChloC^CA^ and Aurora^CA^) except for iChloC, which had a slightly elevated light sensitivity compared to its closing time constant (Figure 2G). Aurora (*ED*_50_ = 0.037 ± 0.003 mW/mm^2^; n = 5) already showed a higher light sensitivity compared to Phobos and iC++ (*ED*_50_ = 0.22 ± 0.06 mW/mm 2, n = 5 and 0.21 ± 0.01 mW/mm2, n = 6) due to slower off kinetics (Figures 2A-C, F, S4). Furthermore, Aurora and Aurora^CA^ displayed inactivation at high green light intensities (Figures 2B, S4E) as already seen in the action spectra (Figure 1C), indicating secondary photochemistry at high light intensities^34^. Orange light (590 to 600 nm) could be used to excite Aurora at the second peak (Figure 1C). However, light sensitivity was 14 fold lower compared to 530 nm, but only 2-3 fold lower compared to Phobos or iC++. Upon activation with 490 nm light iC++^CA^ partially inactivated already at medium light intensities, but blue-shifting the activation wavelength abolished partial inactivation (Figures 2B, S4C). The oƒƒ-kinetics of step-function eACRs could be accelerated by light in a wavelength and intensity-dependent manner as reported for other step-function ChR variants^4,19,20,35^. To determine their inactivation spectra, we activated the step-function eACRs (10 ms) and applied a second light pulse of varying wavelengths but equal photon irradiance (Figure 2D). The maximal inactivation of slow-cycling eACRs was found at 589 nm (n = 6, iC++^CA^/SwiChR++), 603 nm (n = 5, iChloC^CA^), 580 nm (n = 4, Phobos^CA^) and 626 nm (n = 5, Aurora^CA^) (Figure 2E). Maximal additional activation was slightly blue-shifted compared to the respective parental fast-cycling eACR (Figure 2E).

Full channel closing of slow-cycling eACR variants was achieved within seconds or less with light red-shifted to the activation light. For Phobos^CA^ the closing kinetics were accelerated 3 orders of magnitude from 249 ± 10 s (n = 4) to 391 ± 15 ms (n = 5) with 590 nm light. Closing time constant of the iC++ C128A mutation was 115 ± 9 s (n = 7) and could be accelerated 3 orders of magnitude by 600 nm light to 150 ± 15 ms (n = 7), while the Aurora step-function variant C128A could be accelerated from 424 ± 15 s to 916 ± 43 ms (n = 5) with 635 nm light (Figure 2F). As light-accelerated closing is a function of light energy, the *off*-kinetics of eACR C128A variants might be further accelerated with more intense red light^35^. We tested this for iChloC^CA^, where closing could be accelerated from 128 ± 9 s (n = 5) to 346 ± 20 ms (n = 8) at the maximal intensity available (1.95 mW/mm^2^) at 610 nm (Figures 2F, S4F, G). Taken together, optical stimulation of step-function eACRs with light protocols of variable intensities and color, allows temporally precise toggling between open and closed states (Figure 2H).

Next, we systematically compared the following additional parameters of Phobos, Aurora, iC++, iChloC and their respective C128A variants in HEK cells: photocurrent amplitude, reversal potential and membrane targeting. First, to determine membrane localization with higher precision than possible with our initial conventional epifluorescence images (Figure S2, S3), we imaged mCherry labeled eACRs, in HEK cells labeled with the membrane marker Vybrant® DiO by confocal microscopy. All eACRs showed almost exclusive plasma membrane localization with no or only minor fractions of protein found in intracellular compartments (Figure 3A). The relative membrane targeting was >89 % (n = 11 to 21) for all constructs (Figure 3B). At saturating light intensities and wavelengths (*c.f*. Figure 1D, E) average stationary photocurrents of fast-cycling eACRs ranged between 560 and 720 pA, similar to iC++ and two times larger than iChloC. Except for iChloC, all slow-cycling eACR variants had 40 to 60 % smaller current amplitudes compared to their parental constructs (Figure 3C) caused by photochemical back-reactions. When activated with 590 nm, which corresponds to the second activation peak, photocurrents of Aurora and Aurora^CA^ reached 69% of the green light evoked photocurrent. Finally, we verified chloride selectivity by measuring the reversal potential (*E*_rev_) for light-evoked photocurrents under asymmetrical Cl^−^. *E*_rev_ deviated only minimally from the Cl^−^-Nernst potential under the measured conditions for all eACRs (-65.1 ± 0.3 mV, n = 5 to 8) confirming their high chloride selectivity (Figure 3D).

**Figure 3:**
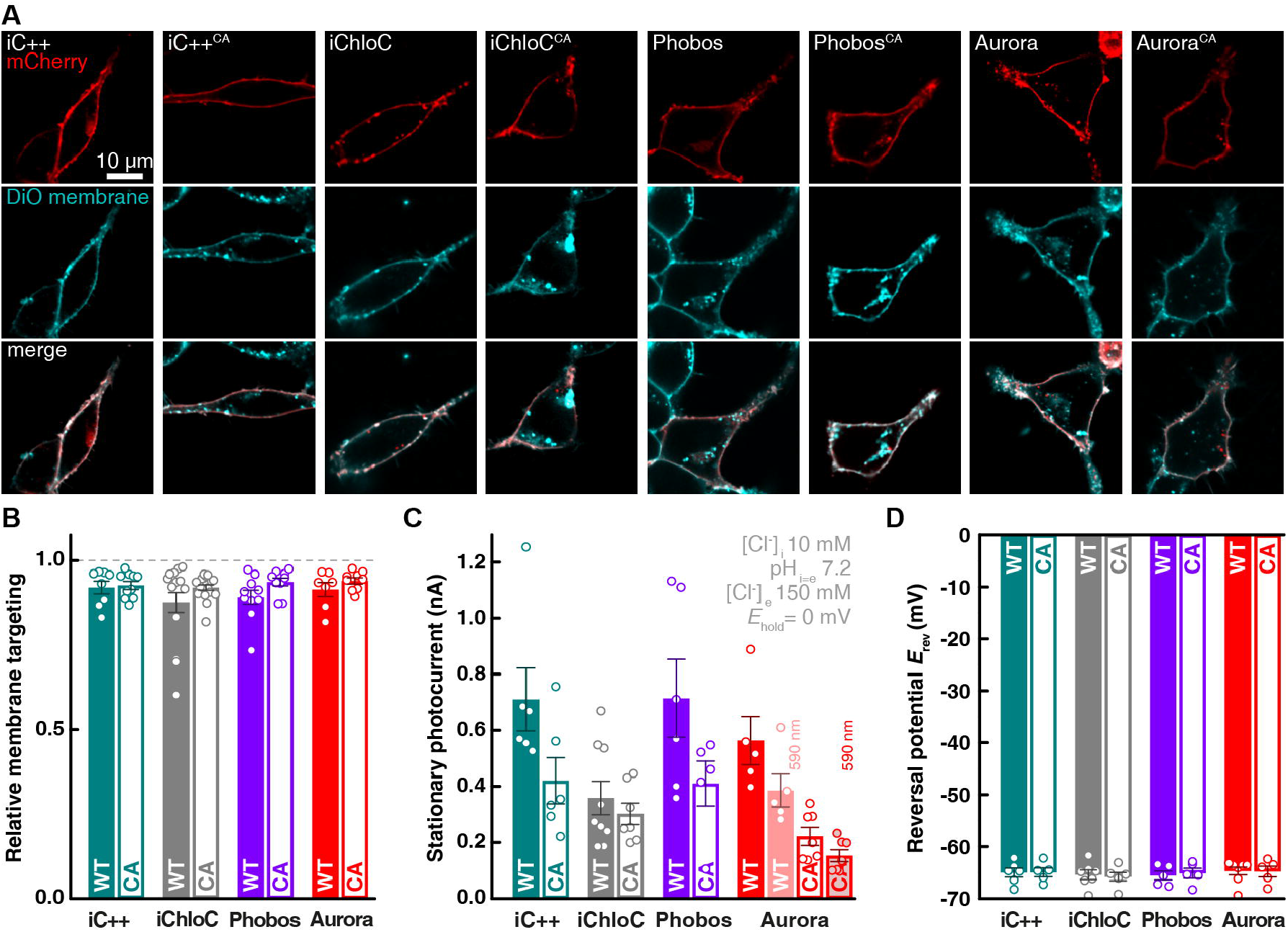
Membrane targeting, amplitudes and reversal potentials of eACRs. **(A)** HEK cells expressing the established eACRs iC++, iChloC and their C128A variants or the newly generated eACR Phobos and Aurora and their C128A variants fused to the mCherry were co-labelled with the membrane dye Vybrant®-DiO (middle row). Upper row: confocal images of mCherry, lower row: merged images of eACR-mCherry and labelled cell membrane (equatorial z-slices). Intensity and/or contrast for DiO were adjusted due to different staining efficiency. **(B)** Relative membrane targeting of eACRs (n = 14 Phobos, 12 Phobos^CA^, 13 iC++, 13 iC++^CA^, 11 Aurora, 12 Aurora^CA^, 18 iChloC, 21 iChloC^CA^). **(C)** Absolute stationary photocurrents of eACRs at indicated conditions (n = 6 Phobos, 5 Phobos^CA^, 6 iC++, 6 iC++^CA^, 5 Aurora, 7 Aurora^CA^, 9 iChloC, 7 iChloC^CA^). **(D)** Reversal potentials (*E*_rev_) for all eACRs at high-extracellular (150 mM) and low-intracellular (10 mM) chloride concentrations (n = 5 Phobos, 5 Phobos^CA^, 6 iC++, 5 iC++^CA^, 8 Aurora, 5 Aurora^CA^, 7 iChloC, 6 iChloC^CA^). Bar plots show mean ± SEM. Single measurement data points are shown as dots.

### Photocurrents and spike inhibition in hippocampal neurons

After biophysical characterization of Aurora and Phobos including their respective step-function variants in HEK cells, we tested their performance in hippocampal neurons. Citrine-labeled Aurora or Phobos was co-expressed with mCerulean in CA1 pyramidal neurons of organotypic hippocampal slice cultures. Four to five days after single-cell electroporation, transfected neurons could be readily identified by their volume marker mCerulean. CA1 neurons had normal morphology and showed bright, membrane-localized expression of the Citrine-labeled eACRs, indicating proper membrane insertion of the eACRs (Figure 4A, E). Notably, in some cases, strong overexpression of Citrine-labeled eACRs yielded additional bright puncta indicative of protein mislocalization and intracellular accumulation^36^ (Figure 4A). However, measuring basic biophysical membrane properties of eACR-expressing CA1 neurons in the dark, using whole-cell patch-clamp experiments, confirmed that overexpression of Phobos or Aurora did not have detectable effects on neuronal function (Supplementary table 1). We next measured the action spectra of Aurora and Phobos with continuous illumination to assess their utility as color-shifted silencing tools in neurons. CA1 neurons were voltage-clamped at −50 mV, approx. 25 mV above the calculated Nernst potential for chloride (-75.9 mV). Under these conditions, entry of Cl^−^ ions resulted in outward-directed photocurrents. All experiments were done in the presence of blockers of ionotropic synaptic transmission. Similar to HEK cell experiments, the action spectrum of Aurora was red-shifted and photocurrents showed a fast inactivating component between 470 nm and 550 nm (Figure 4B, S5). Illumination with orange-red light produced photocurrents with a slower onset and lack of the fast component due to reduced absorption cross-section at this wavelength. Tonic photocurrents still reached 57 ± 3 % of the maximal stationary current at 595 nm and 34 ± 4 % at 635 nm. Thus, orange-to-red light is suitable to evoke photocurrents in Aurora-expressing neurons (Figure 4B, S5). To test the ability of Aurora to block action potentials at various wavelengths, current-clamped neurons were depolarized by somatic injection of 500 pA for 500 ms, triggering typically 7-15 action potentials (Figure 4C). During current injection, light pulses (200 ms) ranging from 365 nm to 660 nm at intensities of 0.1, 1 and 10 mW/mm^2^ were applied. Even at the lowest light intensity (0.1 mW/mm^2^), >80 % of action potentials were blocked between 470 and 525 nm, which is in accordance with our HEK cell measurements of Aurora’s action spectrum (Figure 4D). Under high-intensity illumination (10 mW/mm^2^), however, the spectral range for efficient blocking (>80%) was largely extended, reaching from 365 to 595 nm (Figure 4C, D). Thus, if a high-intensity light source is available, excitation at the edge of Aurora’s action spectrum (595 nm, orange) is sufficient to block action potentials in pyramidal cells.

**Figure 4:**
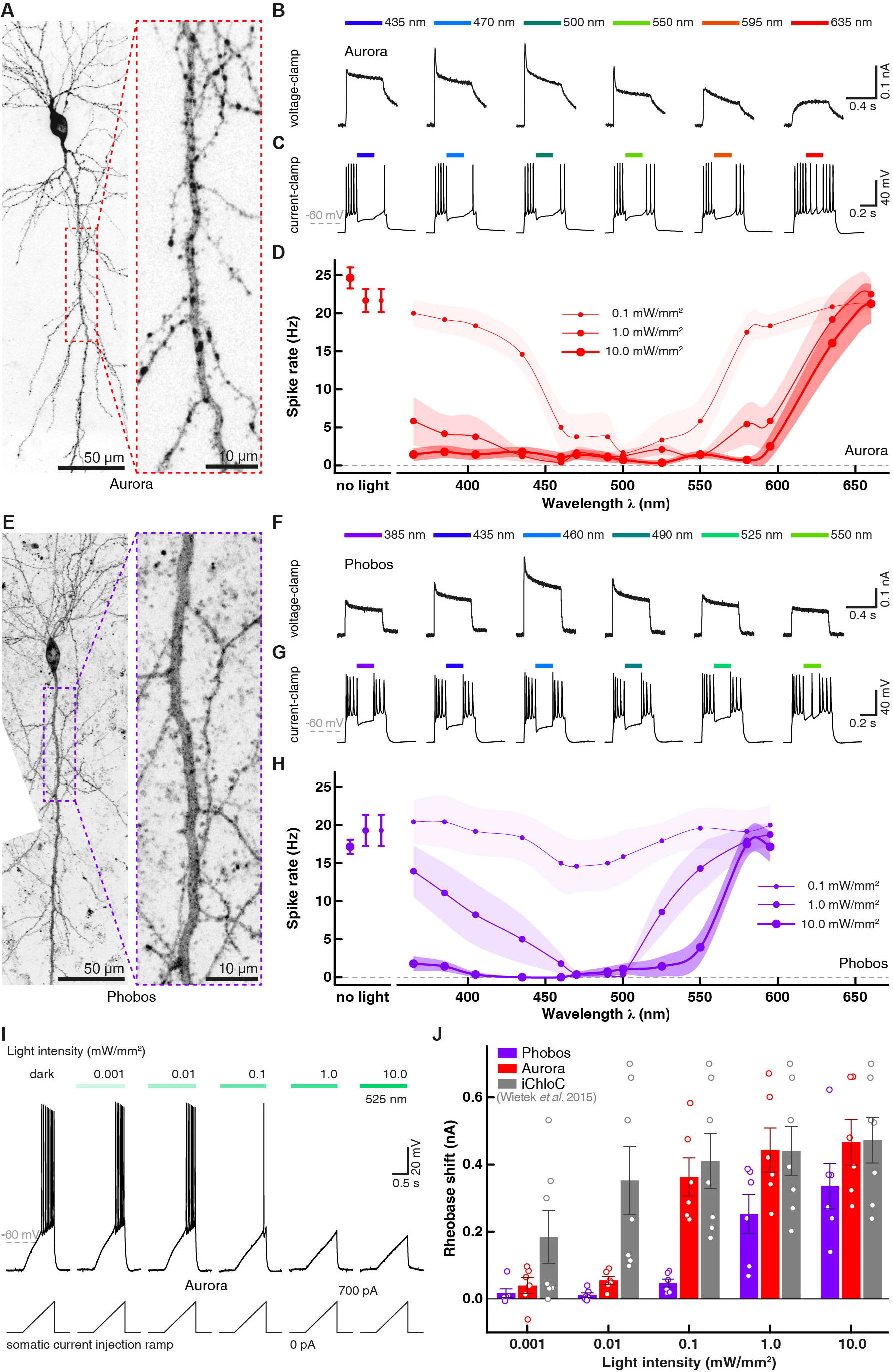
Phobos and Aurora in CA1 pyramidal cells in organotypic hippocampal slice culture. **(A)** CA1 pyramidal neuron expressing Aurora-Citrine 5 days after electroporation (stitched maximum intensity projections of two-photon images, fluorescence intensity shown as inverted gray values). Citrine fluorescence was mainly localized to the plasma membrane across the entire cell. Inset shows magnified view of the apical dendrite. Punctate structures probably arise from protein aggregates. **(B)** Representative photocurrent traces evoked at indicated wavelengths (10 mW/mm^2^). **(C)** Membrane voltage traces in response to 500 ms current injections and 200 ms light pulses at indicated wavelengths (10 mW/mm^2^). Light application was delayed by 144 ms with respect to current onset. **(D)** Quantification of action potential inhibition at indicated light intensities and wavelengths (n = 6 to 7). Lines are interpolations of data points and shaded areas represent SEM. (**E** to **H**) same as (A to D) for Phobos-Citrine expressing neurons (H, n = 4 to 7). **(I)** Voltage traces in response to depolarizing current ramps (0-700 pA) injected into an Aurora-expressing CA1 pyramidal cell. The injected current at the time of the first spike was defined as the rheobase. Illumination with green light activated Aurora. Increasing light intensities shifted the rheobase to higher values. **(J)** Quantification of the rheobase shift generated by Phobos or Aurora compared to iChloC^6^ during illumination with increasing light intensities. Bar plots show mean ± SEM. Single measurement data points are shown as dots.

Consistent with our measurements in HEK cells, Phobos showed a blue-shifted action spectrum in neurons, which peaked at 460 nm and was truncated in the long-wavelength light regime (Figure S5). The peak current was reduced to half-maximum between 500 and 525 nm (500 nm: 68.8 ± 5 % and 525 nm: 41 ± 5 %). Like Aurora, Phobos generated a fast phasic current component, which was absent when activated with more red-shifted light (Figure 4F). As for Aurora, the ability to block action potentials was tested with illumination at various wavelengths (365 to 635 nm) and intensities (0.1, 1, and 10 mW/mm^2^) for 200 ms during 500 ms depolarization steps. In agreement with the measured action spectrum, spikes were efficiently blocked (>90%) with light up to 525 nm at a saturating intensity of 10 mW/mm^2^. At 1 mW/mm^2^ spike block was only efficiently achieved between 435 to 500 nm. With 0.1 mW/mm^2^ action potentials were not efficiently blocked anymore. The maximal effect (50 %) was reached at 460 nm, the peak of the action spectrum of Phobos (Figure 4G, H, S5).

To assess the silencing performance of Aurora and Phobos at their optimal activation wavelengths more quantitatively, we measured their ability to shift the rheobase upon activation at different light intensities. Depolarizing current ramps (from 0-100 to 0-700 pA) were injected into the somata of Aurora - and Phobos-expressing neurons in the dark and during illumination with green (525 nm) or blue (460 nm) light respectively, at intensities ranging from 0.001 to 10 mW/mm^2^ (Figure 4I). A significant shift of the rheobase towards larger currents was detected in Aurora-expressing neurons during illumination at intensities starting from 0.1 mW/mm^2^ (p= 0.04, one-way ANOVA followed by Dunnett’s multiple comparison test n= 6, Figure 4J), while Phobos-expressing cells required illumination with at least 1 mW/mm^2^ for a significant rheobase shift (p= 0.009, one-way ANOVA followed by Dunnett’s multiple comparison test, n= 6, Figure 4J), reflecting a higher light sensitivity of Aurora expressing cells (Figure 2G). Comparing the two novel eACRs to iChloC^6^, the maximal rheobase shifts were similar, indicating that all three eACRs have the same silencing efficacy at high light intensities. However, in agreement with the relationship between closing time constant and relative light sensitivity (Figure 2G), fast-cycling eACRs require more light to reach their maximal silencing capacity.

Next we asked whether the bi-stable variants of Phobos, Aurora and iChloC (C128A) are suitable to block action potentials for an extended time period after a brief light flash and whether this block could be reverted with red-shifted illumination, as suggested by the HEK cell measurements (Figure 2H). Like their parental constructs, Phobos^CA^, Aurora^CA^ and iChloC^CA^ were fused with Citrine and expressed together with mCerulean (Figure 5D, S6A-B). All three step-function eACRs produced photocurrents in voltage clamp experiments. Similar to HEK-cell measurements the net activation spectra were slightly blue-shifted compared to the parental fast cycling eACRs. Also, inactivation spectra peaked at similar wavelengths (Figures 5A-C, S6C-F). To assess spike-block performance, we repeatedly injected, depolarizing current steps (2 s duration) at an interval of 0.2 Hz in current clamp for one minute, which reliably evoked action potential firing (Figure 5E, S7). To open the respective step-function eACR and thereby inhibit action potentials, we applied a 20 ms light flash (Phobos^CA^: 460 nm, iChloC^CA^: 470 nm, Aurora^CA^: 525 nm) after 5 s. Action potentials were efficiently blocked by Phobos^CA^ and iChloC^CA^ in the following 55 s (Figure S7). Also activation of Aurora^CA^ resulted in long-lasting inhibition of action potential firing. However, we noted a slight depolarization of the resting membrane potential in Aurora^CA^ expressing cells during current injection in the dark (Figure S7A) indicating a depolarizing leak conductance in Aurora^CA^ expressing neurons, which is further supported by altered membrane parameters and spike properties in the dark (Supplementary table 1). Moreover, the membrane markedly depolarized after light activation, which prevented complete spike block. These limitations have to be taken into account when considering Aurora^CA^ as a silencing tool in neurons.

**Figure 5:**
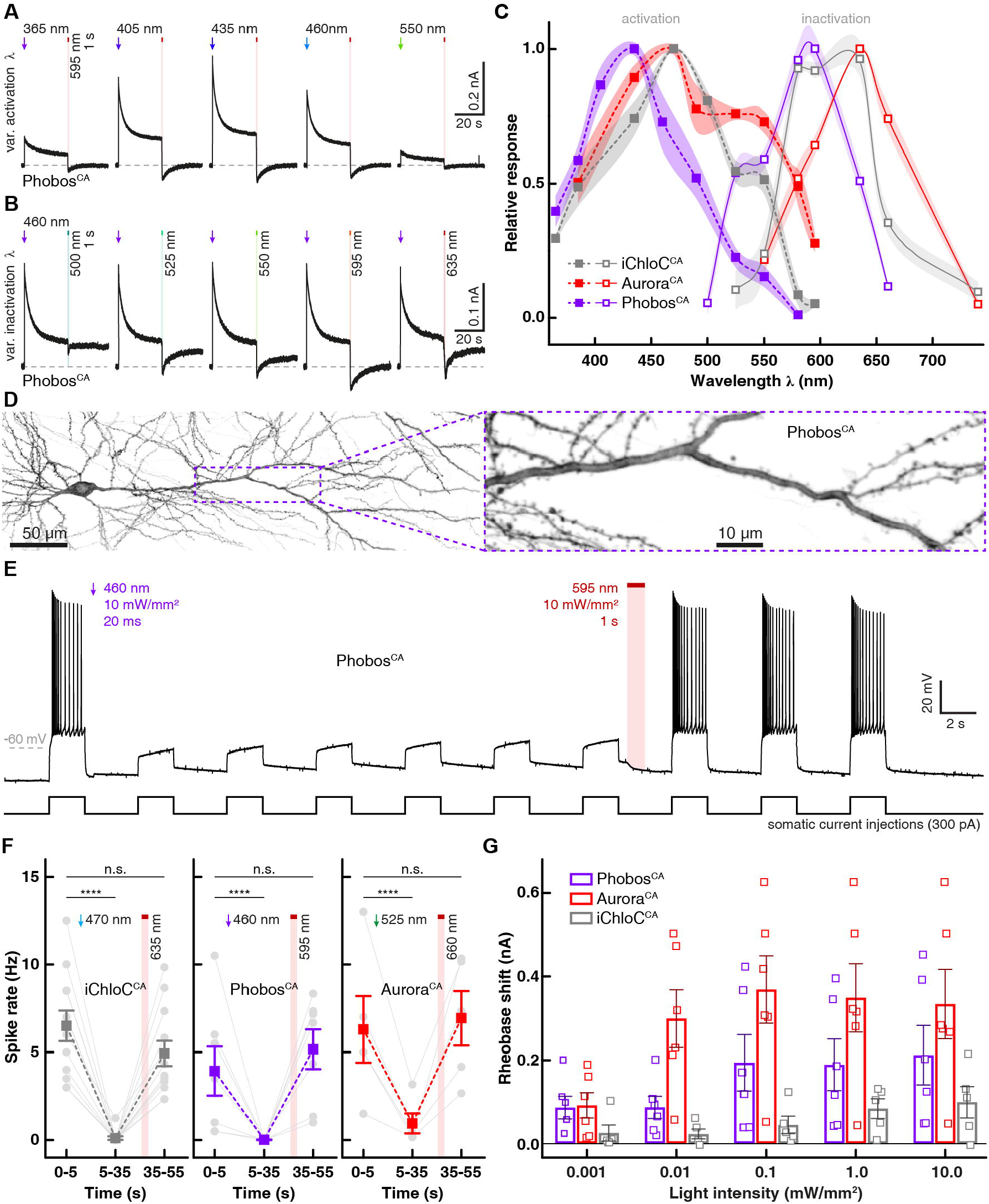
iChloC^CA^, Phobos^CA^ and Aurora^CA^ in CA1 pyramidal cells in organotypic hippocampal slice cultures. **(A)** Representative photocurrent traces of a Phobos^CA^ expressing CA1 cell evoked with different activation wavelengths and shutoff with 595 nm light. **(B)** Photocurrent traces in the same cell evoked with 460 nm light and shutoff with indicated wavelengths (10 mW/mm^2^). **(C)** Activation spectra (dashed lines) and inactivation spectra (solid lines) of Phobos^CA^, iChloC^CA^ and Aurora^CA^ in CA1 pyramidal neurons. Lines are interpolations of data points and shaded areas represent SEM (n = 2 to 9). **(D)** CA1 pyramidal neuron expressing Phobos^CA^-Citrine 5 days after electroporation (stitched maximum intensity projections of two-photon images, fluorescence intensity shown as inverted gray values). Citrine fluorescence was mainly localized to the plasma membrane across the entire cell. Inset shows magnified view of the apical dendrite. **(E)** Membrane voltage trace shows reversible suppression of depolarization-induced spiking by photoswitching Phobos^CA^ between open and closed state. **(F)** Quantification of the spike rate during current injection at indicated time intervals before opening light pulse, after opening light pulse and after closing light pulse in CA1 neurons expressing iChloC^CA^-Citrine (n = 12 neurons in 12 slice cultures), Phobos^CA^-Citrine (n = 7) or Aurora^CA^-Citrine (n = 5). Gray symbols indicate individual experiments. Mean values are shown as rectangular symbols with SEM. ****: p < 0.0001, repeated measures one-way ANOVA followed by Tukey’s multiple comparisons test. **(G)** Quantification of the rheobase shift generated by Phobos^CA^ or Aurora^CA^ and iChloC^CA^ during illumination with increasing light intensities. Bar plots show mean ± SEM. Single measurement data points are shown as open squares.

In a subset of experiments, eACRs were closed with a second light flash (1 s, Phobos^CA^: 595 nm, iChloC^CA^: 635 nm, Aurora^CA^: 660 nm) 30 s after the first light flash. Action potentials immediately returned to the same frequency as before the first light flash, indicating complete shut-down of the chloride conductance (Figure 5E, F). Since acceleration of channel closing depends on the light energy absorbed by the open channel (Figure S4F, G), a longer illumination period at lower light intensities can be used if light power is limited (Figure S6G-I).

While action potentials, which were evoked by depolarizing the cells to potentials just above the firing threshold, were efficiently silenced with all step-function-eACRs, we sought to compare their performance more quantitatively. Rheobase measurements under illumination with blue (460 nm for Phobos^CA^, 470 nm for iChloC^CA^) or green (525 nm for Aurora^CA^) light revealed high silencing efficacy for Aurora^CA^ (maximal rheobase shift: 379 ± 74 pA, n= 6), while iChloC^CA^ shifted the rheobase only marginally (maximal rheobase shift: 99 ± 34 pA, n= 5). Phobos^CA^ shifted the rheobase by 212 ± 65 pA (n= 6, Figure 5G). The slow closing time constant of all three step-function eACRs suggested high operational light sensitivity in neurons (Figure 2G). Indeed, illumination of Aurora^CA^ already at 0.01 mW/mm^2^ was sufficient to elicit a significant rheobase shift (p= 0.02). However, due to the smaller photocurrents of Phobos^CA^ and iChloC^CA^, higher light intensities were required for a significant silencing effect (Phobos^CA^ >0.1 mW/mm^2^, p= 0.006, iChloC^CA^ >1 mW/mm^2^, p= 0.003, Figure 5G). Taking into account the toxic effects of Aurora^CA^ expression and the weak performance of iChloC^CA^, we considered Phobos^CA^ the most useful step-function eACR to silence neurons *in vivo* and focused our efforts on Phobos^CA^ together with the fast cycling Aurora and Phobos variants.

### Light modulation of behavioral responses in *Drosophila*

To test whether eACRs can functionally inhibit neurons *in vivo*, we used *Drosophila melanogaster* as a model organism. We tested whether the novel eACRs can functionally inhibit the larval nociceptive and motor systems. First, we compared the capability of previously published enhanced halorhodopsin eNpHR^37^ with iChloC^6^ and Aurora to inhibit nociceptive class IV da (C4da) neurons, which mediate larval nocifensive rolling responses to mechanical stimulation^38,39^. All animals were raised in the presence of all*-trans* retinal (ATR). eNpHR expression in C4da neurons caused only unspecific defects in nociceptive responses and light-activation of eNpHR had no significant further effect (Figure 6A). In contrast, iChloC activation by blue (460-495 nm) or green-yellow (545-580 nm) light significantly reduced mechano-nociceptive responses. Notably, blue but not green-yellow light illumination alone increased nociceptive responses in control animals that did not express light-activated channels, likely due to the innate blue light sensitivity of C4da neurons^17^. Activation of Aurora with 545-580 nm light strongly reduced nociceptive rolling, suggesting efficient silencing of C4da neurons at a wavelength that does not facilitate nociceptive responses (Figure 6A).

**Figure 6:**
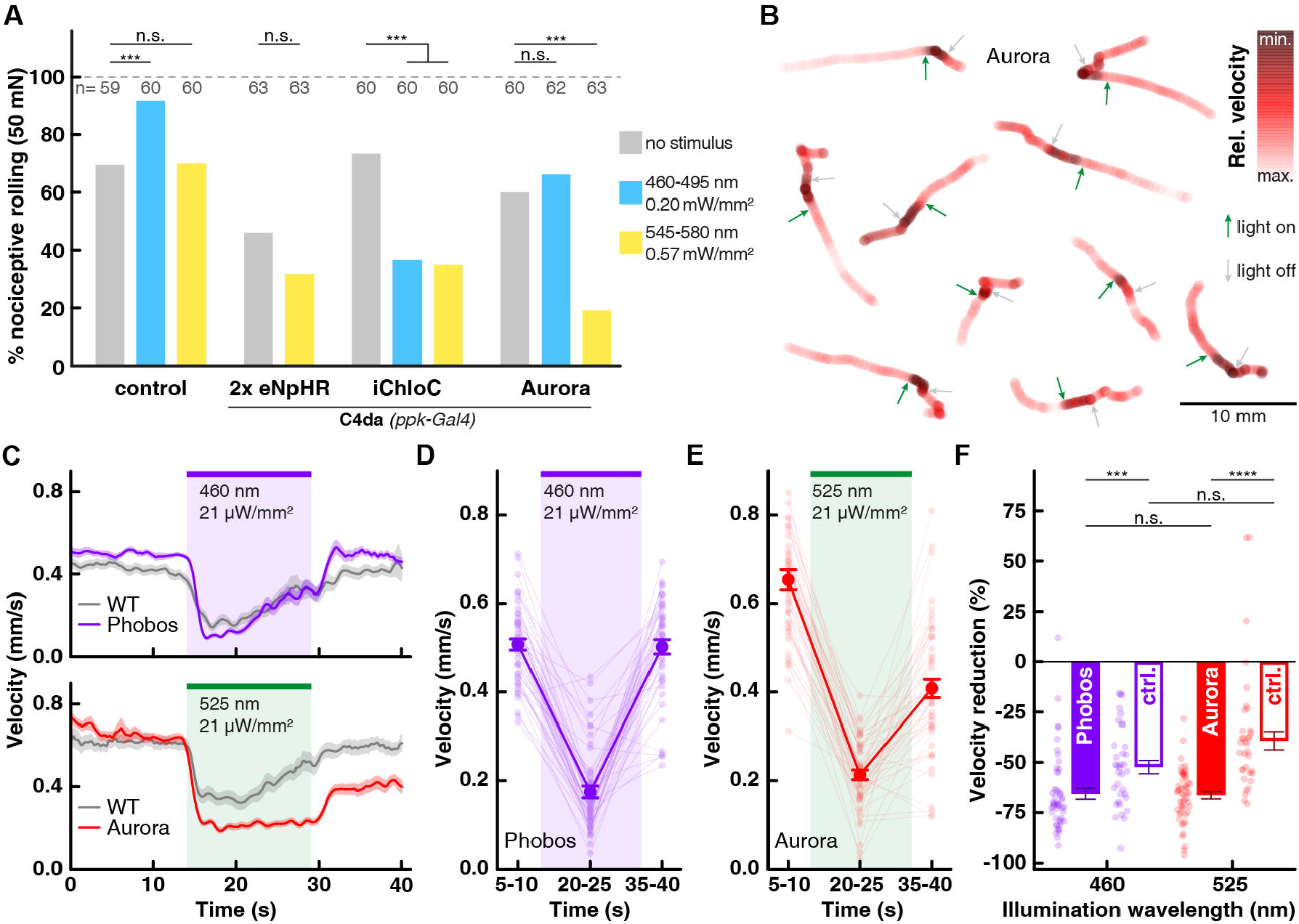
eACRs in *Drosophila* larval nociception and locomotion. **(A)** Mechano-nociceptive responses (rolling) of 3^rd^ instar larvae after 50 mN stimulation with a *von Frey* filament, with and without light activation of C4da neurons (*ppk-Gal4*) expressing Halorhodopsin (2x eNpHR), iChloC or Aurora. No light stimulus (gray), 460-495 nm (blue) and 545-580 nm (yellow) conditions are shown as indicated by color (n as indicated, ***: p< 0.001, chi^2^ test). **(B)** Representative traces of freely locomoting larvae expressing Aurora in motor neurons (*ok371-Gal4*). Arrows indicate onset (green, 525 nm, 21 μW/mm^2^) and offset (gray) of light activation. Relative velocity is color intensity coded in red. **(C)** Average larval velocity over time is plotted for Phobos (top) or Aurora (bottom) expressing animals and respective wild-type controls (*ok371-Gal4*) with activation using a 460 nm or 525 nm light pulse for 15 s, respectively (n = 49 animals for Phobos, n = 52 animals for Aurora, mean ± SEM). (**D** and **E**) Averaged velocity before (5-10 s), during (20-25 s) and after (35-40 s) light induced activation of **(D)** Phobos or **(E)** Aurora (n = 49 animals for Phobos, n = 52 animals for Aurora, mean ± SEM). **(F)** Relative velocity reduction during light for Phobos, Aurora and control animals using 460 nm or 525 nm light, respectively (mean ± SEM, ***: p < 0.001, ****: p < 0.0001, non-parametric Kruskal-Wallis test followed by Dunn’s multiple comparisons test).

We next compared the functionality of the newly generated eACRs in larval locomotion. We expressed eACRs in larval motor neurons and first analyzed their expression and localization. Aurora, Phobos and Phobos^CA^ were homogenously expressed in motor neurons, localizing to dendrites, axons and neuromuscular junctions (NMJs). All three eACRs colocalized with a cell surface marker suggesting efficient surface delivery (Figure S8). Importantly, NMJ morphology was not affected by eACR overexpression, indicating high tolerance in *Drosophila* neurons. We next assessed their efficiency in inhibiting locomotion upon light stimulation. Both, Phobos and Aurora-expressing animals that were raised in the presence of ATR slowed down significantly during a 15 s illumination period by 67.4 ± 2.6 % and 66.3 ± 1.8%, respectively (p< 0.0001, n= 57 and p< 0.0001, n= 52 animals respectively, repeated measures one-way ANOVA, followed by Sidak’s multiple comparisons test). After the light stimulus, the animals accelerated again (Figure 6B-E). However, Phobos expressing animals were already recovering locomotion during light stimulation, suggesting insufficient silencing or desensitization of the channel. In contrast, Aurora expressing animals only accelerated after the illumination period. Their apparent incomplete recovery to full locomotion velocity after light stimulation was partly due to reorientation and avoidance of the edges of the arena (Video 4). Due to the strong innate behavioral response to visible light^17^, wild-type control animals also significantly reduced their velocity upon illumination (470 nm: 52.3 ± 4.5 %, p< 0.0001, n= 35; 525 nm 39.2 ± 4.5 %, p< 0.0001, n= 34; Figure 6F, S9A). However, this reduction was significantly smaller than in animals expressing Aurora or Phobos (Figure 6F). Importantly, wild-type animals were not motionless during illumination. They rather displayed stereotypic head-turns, which signify the innate escape response and therefore reduced linear locomotion speed^40^ (Videos 1&2). In contrast, activation of Phobos or Aurora also abolished head-turning and therefore efficiently inhibited the motor system (Videos 3&4).

Due to the direct impact of continuous illumination on locomotion, we reasoned that the step function mutations should allow us to uncouple the innate light response from the neuronal silencing effect. We found that Aurora^CA^ expression in larval motor neurons was toxic, perhaps due to leak currents in the dark. In contrast, animals expressing iChloC^CA^, were normal, but locomotion could not be inhibited with light (not shown), probably due to the low rheobase shift achieved with iChloC^CA^ (Figure 5G, S6B). Phobos^CA^, on the other hand, was well tolerated by larvae, and activation was able to fully inhibit larval locomotion, which was sustained for at least 120 s after the light pulse (p< 0.0001, n= 52 animals, repeated measures one-way ANOVA, followed by Sidak’s multiple comparisons test, Figure 7B, Video 5). Spontaneous recovery of coordinated forward locomotion occurred at slightly different time points in individual animals (indicated by asterisks in video 5) and was often not observed during the entire recording period of 210 s (Figure 7A, B), suggesting a long-lasting silencing effect of Phobos^CA^ on the motor system of *Drosophila* larvae. However, reverting the open-state of Phobos^CA^ with illumination at 595 nm efficiently recovered larval locomotion during the following 20 s (Figure 7C, D, Video 6), suggesting that the long-lasting inhibition of locomotion was due to Phobos^CA^ conductance. In contrast, wild-type control animals only slowed down during the blue light pulse and regained full locomotion speed after light shutoff (Figure S9B, C, Video 7). These results show that step-function eACRs can be used to modulate *Drosophila* behavior in a binary manner.

**Fig. 7:**
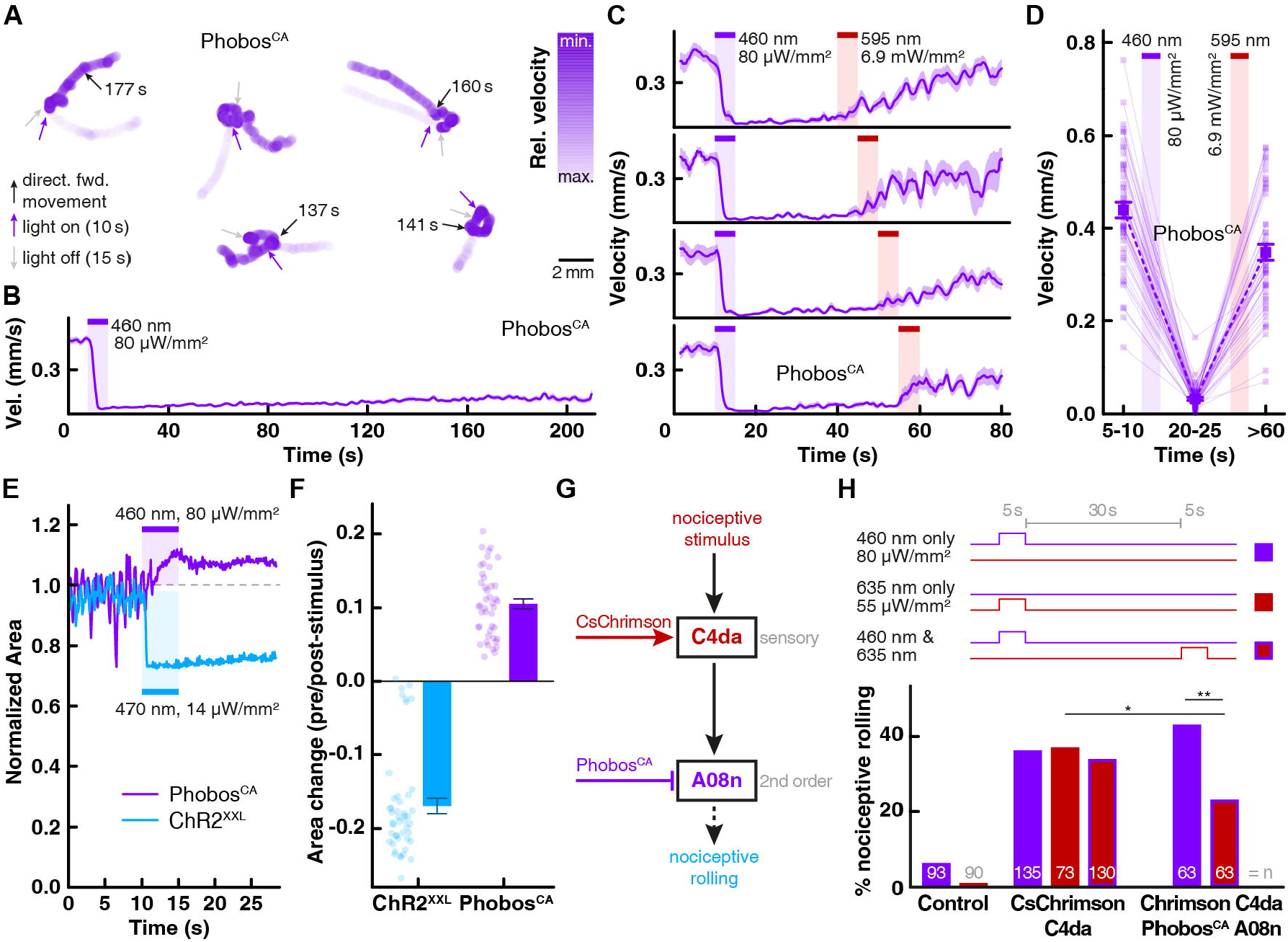
Phobos^CA^ in *Drosophila* larval locomotion and nociception. **(A)** Representative traces of freely locomoting larvae expressing Phobos^CA^ in motor neurons (*ok371-Gal4*). Blue and gray arrows indicate onset and offset of 5 sec blue light pulse, respectively (460 nm, 80 μW/mm^2^). Black arrows indicate time point when animals spontaneously regained coordinated forward locomotion. Relative velocity is color intensity coded in blue. **(B)** Average larval velocity over time (n= 52 animals). **(C)** Average larval velocity over time showing inhibition of locomotion after light induced Phobos^CA^ activation (460 nm, 80 μW/mm^2^) and recovery by channel closing with a 5 s 595 nm light (6.9 mW/mm^2^) at indicated time points (40-60 s). **(D)** Average velocities before (5-10 s) light induced activation of Phobos^CA^, during inhibition (20-25 s) and after channel closing with red-shifted 595 nm light (>60 s) (n= 54 animals, mean ± SEM). **(E)** Comparison of relative larval body size change after ChR2^XXL^ mediated activation of motor neurons and Phobos^CA^ mediated inhibition. Normalized larval body area is plotted over time with indicated light activation of Phobos^CA^ or ChR2^XXL^. **(F)** Quantitative comparison of the relative area change before and after light mediated activation of ChR2^XXL^ or Phobos^CA^ (n= 44 animals for Phobos^CA^, n= 46 animals for ChR2^XXL^, mean ± SEM). **(G)** Simplified scheme of the mechano-nociceptive circuit in *Drosophila* larvae. Red-light-activation of CsChrimson-expressing C4da neurons triggers nociceptive rolling^39^. Prior blue-light activation of Phobos^CA^ in A08n neurons downstream of C4da neurons is expected to reduce CsChrimson-evoked nociceptive rolling. **(H)** Light-induced mechano-nociceptive responses (rolling) of 3^rd^ instar larvae expressing CsChrimson in C4da neurons (*27H06-LexA*) alone or together with Phobos^CA^ in A08n neurons (*82E12-Gal4*). Wild-type larvae serve as controls. Light stimulus conditions are shown as indicated by color (n as indicated, *: p< 0.05, **: p< 0.01, chi^2^ test).

To confirm that eACRs indeed have an inhibitory silencing effect on the motor system, we compared the effect of Phobos^CA^ activation on larval body length with that of Channelrhodopsin-2-XXL (ChR2^XXL^)^41^. ChR2^XXL^ activation in motor neurons resulted in strong body wall muscle contraction and an effective shrinkage of the detectable larval body area, consistent with motor neuron activation. Conversely, Phobos^CA^ activation resulted in body wall muscle relaxation indicated by an increase in the detectable area due to lack of contracting segments (Figure 7E, F), confirming the inhibitory nature of Phobos^CA^ action.

Finally, to provide proof of the concept that silencing with Phobos^CA^ can be combined with other, spectrally distinct optogenetic tools to bidirectionally modulate circuit function, we asked whether nociceptive rolling, evoked by CsChrimson-mediated activation of sensory C4da neurons, could be reduced by Phobos^CA^-mediated silencing of downstream-A08n neurons^39^ (Figure 7G). As shown previously, chronic silencing of A08n neurons overexpressing the potassium channel subunit Kir2.1 reduced rolling after CsChrimson-mediated C4da activation by approximately 60%^39^. As expected, illumination of animals expressing only CsChrimson in C4da neurons with low-intensity red light (635 nm, 5 s, 0.055 mW/mm^2^) induced nociceptive rolling which was not observed in wild-type animals (Figure 7H). However, also blue light, which only marginally elevated rolling in control animals, triggered rolling in CsChrimson animals indistinguishable from red-light activated rolling. The light sensitivity of red-shifted CCRs, including CsChrimson, in the blue spectrum therefore makes exclusive activation of a blue light-sensitive eACR difficult. In addition, the innate blue light sensitivity of *Drosophila* larvae could interfere with the behavioral outcome (Figure 6A). The long-lasting conducting state of Phobos^CA^ allowed us to temporally dissociate the blue-light response from the silencing-effect on CsChrimson-mediated rolling. While blue light to activate Phobos^CA^ triggered acute rolling (probably due to the fast activation of CsChrimson vs. Phobos^CA^), CsChrimson-mediated rolling triggered by a red light pulse 30 s later was significantly reduced (Figure 7H). The red light used to activate CsChrimson (635 nm, 5 s, 0.055 mW/mm^2^) was two orders of magnitude weaker than the orange light required to switch Phobos^CA^ to the closed state (595 nm, 5 s, 6.9 mW/mm^2^), indicating that CsChrimson activation did not directly interfere with Phobos^CA^-function. These results show that Phobos^CA^ can be combined with red-shifted CCRs like CsChrimson to perform functional circuit analysis by activation/inactivation experiments *in vivo*.

## Discussion

We extensively explored the possibilities to engineer novel eACRs from 11 different well-characterized CCRs (Figure S1) by employing two independent strategies based on previous mutagenesis of *Cr*ChR2^7^ or the C1C2 chimera^5^. The first strategy was based on the exchange of an acidic glutamate for a basic arginine at position 90 (E90R) in the central gate of *Cr*ChR2. The second strategy aimed at systematically replacing negative charges in the outer pore of the channel with positive or neutral charges, without compromising the photocycle or protein stability of C1C2. Residual proton conductance of both variants was eliminated in a second round of optimization yielding the two highly chloride selective eACRs iC++ and iChloC^4,6^. The successful conversion of two different, yet closely related CCRs (*Cr*ChR2 and C1C2, figure S10) prompted us to systematically investigate the applicability of the two conversion strategies to other CCRs. We focused mainly on CCRs with blue-or red-shifted action spectra relative to iC++ and iChloC. We also asked if the iC++-strategy was applicable to *Cr*ChR2 and conversely, if the iChloC-strategy was applicable to C1C2. Interestingly, while ion selectivity could be inverted in *Cr*ChR2 with both strategies, conversion of C1C2 with the ChloC approach - albeit successful - yielded low channel expression in HEK cells and negligible photocurrents (Figure S2). Low expression or improper subcellular targeting was also observed for various CCRs converted with both strategies. Thus, both mutagenesis strategies were limited to a small subset of CCRs and not generalizable. The main limitation often appears to be the deleterious effect of mutagenesis on proper folding and plasma membrane localization of the channel. For example, both ChloC and iC++ conversion strategies failed for the three blue-shifted CCRs *Tc*ChR, *Ts*ChR, *Ps*ChR2 and the fast CCR Chronos due to loss of protein expression (except for the ChloC strategy on TcChR, which did not affect protein expression but failed to shift ion selectivity). Similarly, photocurrents of the mutated red-shifted CCRs *Vc*ChR1, C1V1b and Chrimson were extremely weak or absent altogether. However, in this case, despite resulting in poor expression and low photocurrents, the iC++ approach rendered *Vc*ChR1, C1V1b completely anion-selective.

The only CCR where both strategies yielded full anion selectivity without compromising photocurrents except for *Cr*ChR2 was *Co*ChR, which is closely related to C1C2 and *Cr*ChR2 (Figure S10). While the ChloC approach alone resulted in an incomplete shift of the ion selectivity, additional iChloC mutations rendered *Co*ChR completely anion-selective. However, as shown earlier, expression in neurons was cytotoxic, most likely due to leakiness of the channel in the dark^6^. Since the ChloC/iChloC conversion strategy failed in all other CCRs, we assume that the disruption in the central gate caused by the mutation of glutamate to an arginine destabilized the protein. Alternatively, the hydrogen bonding network between amino acids in the central gate may be differently arranged in different CCRs.

Also the iC++ strategy was too disruptive in most cases and only yielded three new eACRs without introducing leakiness or compromising photocurrents. Of these three, two have a similar action spectrum than the original iC++ and only Aurora, which was derived from ReaChR, displayed an action spectrum that was red-shifted compared to existing eACRs. We mapped the CCRs which could be converted with both approaches on a phylogenetic tree. Interestingly, the conversion strategies were only successful in CCRs closely related to *Cr*ChR2 and C1C2 (Figure S10). A blue-shifted eACR could be generated by altering the action spectrum of an existing eACR. This was achieved by converting the two residues T159 and G163 of iC++ to G and A, respectively, similar to the blue/shifted CCRs *Tc*ChR, *Ts*ChR, *Ps*ChR2, yielding Phobos. In summary, we successfully produced both a red-shifted eACR by applying the iC++ strategy to ReaChR and a blue-shifted ACR by introducing two point mutations from blue-shifted CCRs in iC++.

Although these newly generated eACRs were well-tolerated by neurons, as with many rhodopsins^36^, strong overexpression led to cytoplasmic aggregates, suggestive of a residual protein fraction that did not properly integrate into the plasma membrane. Thus, if possible, expression should be tightly controlled and kept at the minimal level required to achieve the desired photocurrents. In the case of iChloC^CA^ we observed strong accumulation in the somatic region, suggestive of misfolded protein aggregates in the endoplasmic reticulum (Figure S6B). This observation may explain the weak photocurrents of iChloC^CA^ in neurons. To improve protein folding and membrane trafficking, addition of endoplasmic and Golgi export signals and a membrane trafficking signal may be considered for future versions of eACR-constructs^36^.

While iChloC^CA^ and Phobos^CA^ did not alter neuronal properties in the dark, Aurora^CA^ apparently was leaky in neurons, resulting in altered membrane properties in hippocampal neurons and developmental problems in *Drosophila*. Thus, despite successful transformation into a step-function eACR, Aurora^CA^ cannot be recommended as an optogenetic tool due its side effects. No such problems were encountered with the original fast-closing version of Aurora. The unexpected red-shift of iChloC^CA^ may be a direct consequence of the C128A and D156N mutations. Both residues are located inside the retinal binding pocket and interact with the retinal Schiff base. The thiol group of C128 was suggested to directly interact with the retinal molecule^33,42^. The altered hydrogen bonding network may lower the absorption energy required to isomerize the bound retinal from *all-trans* to 13-cis.

The photocurrents produced by the natural *Gt*ACR1/2 are higher than those of any other ACR or CCR engineered or discovered so far, probably due to their large unitary conductance^1^. Therefore, nACRs may be favored over eACRs if acute, transient silencing is desired. However, due to their large conductance, illumination of nACRs at the far edges of their absorption spectra may still generate significant photocurrents potent enough to exert neural silencing. Especially during continuous illumination, which is required for neuronal silencing with nACRs, the action spectrum broadens^34^. This limits the combination of nACRs with other light-dependent applications that require illumination with wavelengths close to the borders of the nACR action spectra. Moreover, step-function variants of nACRs that can be toggled between conducting and non-conducting states by light with different wavelengths have not been reported. Engineering true step-function opsins (SFO)^19,20^ from nACRs is not straightforward due to the low degree of homology between nACRs and most CCRs/eACRs^43^. In contrast, the SFO-strategy was previously applied to iC++ where introduction of the C128A mutation yielded the SFO-eACR SwiChR++^4^. In the present study, we demonstrate that Aurora, Phobos and the previously published iChloC can be turned into SFOs by the C128A mutation, slowing down channel closing by up to 4 orders of magnitude. All new eACRs harboring the C128A mutation could be completely closed with light red-shifted by approx. 110 - 170 nm compared to the activation light.

Optogenetic experiments involving behavioral read-out always carry the risk that the activation light is directly sensed by the animal, potentially leading to wrong conclusions about the function of the neural circuit in question. Two strategies may be employed to avoid this experimental problem using SFO-eACRs. First, low-intensity light is integrated over time by SFO-eACRs, accumulating more and more channels in the open state. Compared to inhibition through light-driven ion pumps, where only a single ion is moved per absorbed photon, light-gated channels offer greater efficacy, and SFOs take this principle to the extreme. However, *Drosophila* larvae are detecting and avoiding even low-intensity light^44 45^. Interference with innate responses is avoided by uncoupling functional silencing from the light stimulus. All our new C128A mutants (iChloC^CA^, Phobos^CA^ and Aurora^CA^) are suitable, in principle, to attenuate action potential firing in neurons for at least 50 s after light shutoff (Figure S7C, D). This property enabled us to dissociate the silencing effects of the step-function eACR from the light stimulus in *Drosophila* larvae. Due to the toxicity of Aurora^CA^ and the weak silencing potential of iChloC^CA^, we limited our *in vivo* experiments to Phobos^CA^. Activation of Phobos^CA^ with a brief light pulse stopped locomotion for an extended time period while wild-type larvae showed only a transient response. In addition, locomotion could be restored by illumination with orange light, which does not strongly affect natural behavior^44^. Thus, we provide proof-of-concept that defined neuronal populations in intact animals can be switched *off* and back *on* with brief light pulses, allowing investigation of behavioral effects without continuous illumination.

Our results further show that the diametrically opposed kinetic and spectral properties of Phobos^CA^ and CsChrimson make them a promising pair of tools for the functional dissection of neural circuits. Previous reports highlighted the combined use of channelrhodopsin-and Halorhodopsin-mediated activation and inhibition, respectively^46,47^. However, generalization of these approaches has so far proven difficult due to cross-activation or low efficiency of Halorhodopsin-mediated silencing. A recent study in *Drosophila* used the fast-cycling nACRs *Gt*ACR1 and *Gt*ACR2 to silence neurons, which in principle, can be combination with Chrimson activation^48^. However, due to the fast kinetics of these tools, light has to be applied continuously to inhibit neural populations during simultaneous activation of Chrimson-expressing neurons, making it difficult to interpret the effects of light on behavior. The property of bistable eACRs like Phobos^CA^ to induce prolonged silencing when triggered by a short light pulse circumvents this limitation. Combining optogenetic excitation and optogenetic inhibition of distinct neuronal populations in a single experiment is still challenging, but holds great promise to unravel the function of more complex circuits.

## Methods

### Molecular Biology and eACR variant design

Expression vectors encoding genes for ChRs where constructed using conventional PCR and restriction enzyme-based cloning methods (FastDigest, Thermo Fisher Scientific, Waltham, MA). Briefly, ChR cDNAs were cloned into p-EGFP-C1 vectors using NheI and AgeI restriction sites. EGFP was replaced by mCherry using AgeI and XhoI restriction sites except for iChloC variants where a p-EGFP-N1 vector was used. The QuikChange II kit (Agilent Technologies, Santa Clara, CA) was used to exchange single or multiple amino acids (C128A, T159G and G163A) in ChloC/iChloC based variants. For the iC++ based approach, ChR variants with the replaced N-terminus of iC++ and all iC++ homologous mutations were constructed *in-silico*, synthesized (GenScript, NJ) and cloned into p-EGFP-C1 vectors as described above. Amino acid sequences and mutations can be found in the supplementary material (Figure S1). We deposited the plasmids encoding for eACRs with the Addgene plasmid repository (#98165: p-mCherry-C1-iC++, #98166: p-mCherry-C1-Phobos, #98167: p-mCherry-C1-Aurora, #98168: p-mCherry-C1-iC++_CA, #98169: p-mCherry-C1-Phobos_CA, #98170: p-mCherry-C1-Aurora_CA, # 98171: p-mCherry-N1-iChloC_CA, #98172: p-mCherry-C1-iChR2^T159C^++, #98173: p-mCherry-C1-i*Co*ChR++).

### HEK293 cell culture

HEK-293 cells (ACC-305, catalogue no. 85120602, Sigma-Aldrich, Munich, Germany) were cultured at 5 % CO_2_ and 37 °C in Dulbecco’s minimal essential medium (DMEM) supplemented with 10 % fetal bovine serum (FBS) and 100 μg/ml penicillin/streptomycin (all from Biochrom, Berlin, Germany). Cells where routinely tested with DAPI staining and PCR assay for mycoplasma contamination. For electrical recordings, cells were seeded onto poly-lysine coated glass coverslips at a concentration of 0.5 × 10^5^ cells*ml^−1^ (2 ml total in 35 mm standard cell culture dishes) and supplemented with a final concentration of 1 μM *all-trans* retinal (Sigma-Aldrich, Munich, Germany). For confocal imaging cells were seeded with the same protocol in poly-lysine coated 35 mm glass bottom dishes (MaTek, Ashland, MA). Cells were transfected with eACR-cDNA using Fugene HD (Roche, Mannheim, Germany) 36 h before measurements.

### Electrophysiological recordings in HEK293 cells

eACR expressing HEK cells where patched at low chloride conditions (10 mM intra-and extracellular). In whole-cell configuration the extracellular buffer was changed to high chloride (150 mM), resulting in a liquid junction potential of 10.5 mV that was corrected on-line. The buffer composition for the pipette solution (10 mM Cl^−^) was (in mM): 2 MgCl_2_, 2 CaCl_2_, 1 KCl, 1 CsCl, 10 EGTA, 10 HEPES, 110 Na-Aspartate. The low/high chloride bath solution (10/150 mM Cl^−^) was composed of (in mM): 2 MgCl_2_, 2 CaCl2, 1 KCl, 1 CsCl, 10 HEPES, 0/140 NaCl, 140/0 Na-Aspartate. All buffers were adjusted with N-methyl-D-glucamine to pH 7.2. The final osmolarity was adjusted to 320 mOsm for extracellular solutions and 290 mOsm for intracellular solutions. External buffer solutions were exchanged by perfusion of at least 2.5 ml of the respective buffer into the custom made recording chamber (volume ~500 μl) while the bath level was kept constant with a ringer bath handler (MCPU, Lorenz Messgerätebau, Katlenburg-Lindau, Germany). Patch pipettes were pulled using a P1000 micropipette puller (Sutter Instruments, Novato, CA), and fire-polished. Pipette resistance was 1.5 to 2.5 MΩ. A 140 mM NaCl agar bridge served as reference (bath) electrode. In whole-cell recordings membrane resistance was >500 MΩ (typically >1 GΩ) and access resistance was below 10 MΩ. All experiments were carried out at 25 °C. Signals were amplified (AxoPatch200B), digitized (DigiData1400) and acquired using Clampex 10.4 Software (all from Molecular Devices, Sunnyvale, CA). Holding potentials were varied between −80 and +40 mV as indicated. A detailed protocol can be found in Grimm et. al ^49^.

A Polychrome V light source (TILL Photonics, Planegg, Germany) was used in most HEK-cell experiments. The half band width was set to ±7 nm for all measurements. Actinic light was coupled into an Axiovert 100 microscope (Carl Zeiss, Jena, Germany) and delivered to the sample using a 90/10 beam splitter (Chroma, Bellows Falls, VT). Light exposure was controlled with a programmable shutter system (VS25 and VCM-D1, Vincent Associates, Rochester, NY). Intensities were measured in the sample plane with a calibrated optometer (P9710, Gigahertz Optik, Türkenfeld, Germany). Light intensities were calculated for the illuminated field of the W Plan-Apochromat 40x/1.0 DIC objective (0.066 mm2, Carl Zeiss).

To record action spectra, a motorized neutral density filter wheel (NDF) (Newport, Irvine, CA) was inserted into the light path between the Polychrome V and the microscope to obtain the same photon irradiance for all wavelengths (390 to 670 nm; 10 or 20 nm steps). Custom software written in LabVIEW (National Instruments, Austin, TX) was used for control and synchronization with electrophysiological experiments. Light was applied for 10 ms at 0 mV holding potential. Minimal deviations in photon irradiance were corrected by linear normalization post measurements.

To record inactivation spectra of step-function eACRs, a 150 W Xenon lamp (LOT-QuantumDesign, Darmstad, Germany), filtered with single band pass filters (460±10 nm, 490±8 nm and 520±20 nm) was coupled into the light path using a 30/70 beam splitter (Chroma, Bellows Falls, VT) to activate slow-cycling eACRs. The 10 ms light exposure was controlled with the same programmable shutter system as used for the Polychrome V. Next, light of various wavelengths at the same photon irradiance (adjusted as described above) was applied for 8 s to inactivate (or additionally activate) eACRs, followed by complete channel closing achieved by application of red light for another 8 s. The holding potential was kept at 0 mV.

For light titration experiments, ND filters (SCHOTT, Mainz, Germany) were used for attenuation. Activating light was applied for 1 s (fast cycling variants) 12 s (slow cycling variants). The holding potential was kept at 20 mV. In case of step-function eACRs, channel closing was accelerated with application of red light between single trials (Figure S4A).

### Analysis of HEK293 cell electrophysioiogy

Data were analyzed using Clampfit 10.4 (Molecular Devices, Sunnyvale, CA) and Origin 9 (OriginLab, Northampton, MA). Stationary photocurrents were measured for the last 40 ms of illumination period or for 40 ms, 2 s after activation of step-function variants. To obtain reversal potentials (*E*_rev_), photocurrents were plotted against the respective holding potential. Next, *E*_rev_ was calculated from the intersection of the current-voltage relation with the voltage axis. For action spectra, photocurrents were normalized to the maximum. To determine inactivation spectra (Figure 2E), mean stationary currents before and after additional activation/inactivation light (1 s after light was switched off) were averaged over a 200 ms period. The current difference (before-after) was divided by the current prior to inactivation (Figure 2D) and plotted against the wavelengths. Additional activation was normalized to maximum, whereas inactivation was not further normalized. The maximum response wavelength (λ_max_) was determined by fitting single recorded action spectra with a 3-parameter Weibull function. Half maximal effective light dose values (*EC*_50_) were determined by fitting single light titration curves by logistic growth function. Here, the photocurrents of fast cycling variants were measured for the last 40 ms of the 1 s illumination period. For slow cycling variants photocurrents were evaluated 1 s after beginning of illumination and post illumination (12 s), both averaged over 40 ms (Figure S4A). Kinetic properties were determined by mono-or double-exponential fits and apparent closing constants were reported. For representative closing kinetic traces (Figure 2A), signals were binned to 50 points per decade with a custom written Matlab script (The MathWorks, Natick, MA).

### Epifluorescence and bright-field microscopy

Fluorescence HEK-cell images shown in figures S2 and S3 were acquired with a triple band ECFP/EYFP/mCherry 69008 filter set (Chroma, Bellows Falls, VT, USA) and a Wat-221S CCD camera (Watec, Tsuruoka, Japan) on the same Axiovert 100 microscope (Carl Zeiss) setup used for electrophysiological recordings (see above). Fluorescence excitation of mCherry was performed using Polychrome V set to 590±15 nm. Background-subtracted images were calculated with Fiji^50^.

### Confocal microscopy

Confocal HEK-cell images shown in figure 3 were taken with a FV1000 confocal laser scanning microscope equipped with an UPLSAPO 60XW objective (Olympus, Hamburg, Germany). The membrane of cells expressing eACR variants fused to mCherry was labeled with Vybrant DiO (ThermoFisher Scientific). mCherry was excited with a 559 nm diode laser and DiO was excited with a 488 nm Argon laser. Mean fluorescence intensities (per area) of the respective eACR-mCherry fusion construct either in the cell membrane (identified by DiO) or within the cell were evaluated (mean background fluorescence was subtracted) for three equatorial slices per cell using a custom Fiji macro and then averaged. The relative membrane targeting values were determined by dividing mean fluorescence density in the cell membrane by the sum of the fluorescence densities (membrane and cytosol).

### Two-photon microscopy

Neurons in organotypic slice cultures were imaged with two-photon microscopy (980 nm excitation) to characterize dendritic morphology and the subcellular localization of citrine-labeled eACRs. The custom-built microscope was controlled by ScanImage software (HHMI Janelia Farm)^51^. Green fluorescence was detected through the objective (LUMPLFLN 60XW, Olympus, Hamburg, Germany) and the oil-immersion condenser (1.4 NA) using GaAsP-PMTs (Hamamatsu, Japan).

### Neuronal recordings in hippocampal slice cultures

All eACR mutants were subcloned into identical neuron-specific expression vectors (pAAV backbone, human *synapsin* promoter), followed by the sequence for a citrine fluorescent protein^52^. We deposited the AAV-plasmids encoding eACRs with the Addgene plasmid repository (#98216: Phobos-Citrine, #98217, Aurora-Citrine, #98218: Phobos^CA^-Citrine, #98219: Aurora^CA^-Citrine, # 98220: iChloC^CA^-Citrine). Organotypic slice cultures of rat hippocampus were prepared as described ^53^ and transfected by single-cell electroporation ^54^ after 14 days *in vitro* (DIV). Plasmids were each diluted to 20 ng/μl in K-gluconate-based solution consisting of (in mM): 135 K-gluconate, 4 MgCl_2_, 4 Na_2_-ATP, 0.4 Na-GTP, 10 Na_2_-phosphocreatine, 3 ascorbate, 0.02 Alexa Fluor 594, and 10 HEPES (pH 7.2). An Axoporator 800A (Molecular Devices) was used to deliver 50 hyperpolarizing pulses (-12 mV, 0.5 ms) at 50 Hz. At DIV 18-20, targeted patch-clamp recordings of transfected neurons were performed under visual guidance using a BX-51WI microscope (Olympus), a Multiclamp 700B amplifier (Molecular Devices), and Ephus software (HHMI Janelia Farm) ^55^ or a doubleIPA amplifier (Sutter Instrument) and SutterPatch software (Sutter Instrument). Patch pipettes with a tip resistance of 3-4 MΩ were filled with (in mM): 135 K-gluconate, 4 MgCl_2_, 4 Na_2_-ATP, 0.4 Na-GTP, 10 Na_2_-phosphocreatine, 3 ascorbate, 0.2 EGTA, and 10 HEPES (pH 7.2). Artificial cerebrospinal fluid (ACSF) consisted of (in mM): 135 NaCl, 2.5 KCl, 2 CaCl_2_, 1 MgCl_2_, 10 Na-HEPES, 12.5 D-glucose, 1.25 NaH_2_PO_4_ (pH 7.4). Synaptic currents were blocked with 10 μM CPPene, 10 μM NBQX, and 10 μM bicuculline or 100 μm picrotoxin (Tocris, Bristol, UK). Measurements were corrected for a liquid junction potential of −10.6 mV. A 16-channel pE-4000 LED light engine (CoolLED, Andover, UK) was used for epifluorescence excitation and delivery of light pulses (ranging from 365 to 660 nm). Light intensity was measured in the object plane with a 1918-R power meter equipped with a calibrated 818-ST2-UV/D detector (Newport, Irvine CA) and divided by the illuminated field (0.134 mm^2^) of the LUMPLFLN 60XW objective (Olympus).

### Behavioral assays in Drosophila melanogaster

cDNAs encoding eACR were codon-optimized for *Drosophiia meianogaster*, synthesized (Thermo Fisher Scientific) and cloned into a 20x UAS vector (pJFRC7, Addgene #26220) ^56^ together with a C-terminal mCerulean3^57^ for Aurora, Phobos, and Phobos^CA^, or tdTomato^58^ for iChloC. Transgenic lines were generated in the attP2 locus using phiC31-mediated transgenesis^59^. The following additional lines were used: *UAS-ChR-XXL* ^41^, *UAS-eNpHR3.0-YFP, LexAop-CsChrimson*^25^, *ppk-Gal4^e¤^* and *27H06-LexA*^61^ for C4da, *82E12-Gal4* for A08n^61^, and *vgiuť^k371^-Gai4* ^62^ for motor neuron expression of eACRs.

Embryos from experimental crosses were collected on grape juice agar plates and supplied with fresh yeast paste containing 5 mM *all-trans* retinal (ATR) and kept at 25 °C in the dark. Staged and density controlled 3^rd^ instar larvae (96 h ± 3 h after egg laying) were collected under low red light illumination (>700 nm).

For mechano-nociception, animals were placed on a 2 % agar plate and forward-locomoting larvae were stimulated on mid-abdominal segments (3-5) with a 50 mN *von Frey* filament twice within 2 s. The behavioral response was visually scored under a stereoscope as nonnociceptive (no response, stop, stop and turn) or nociceptive (bending, rolling). For analysis, only nocifensive rolling behavior (full 360° turn along the body axis) was compared. For simultaneous eACR activation, ATR-fed staged and density-controlled 3^rd^ instar larvae (96±3 h after egg laying (AEL)) were exposed to 460-495 nm (0.20 mW/mm2) or 540-580 nm (0.57 mW/mm2) light from a mercury vapor short arc light source under a stereoscope (Olympus SZX16 with X-Cite 120Q illumination system, Excelitas Technologies, Waltham, MA). Each genotype was tested multiple times on different days and data from all trials were combined. Statistical significance was calculated using a chi^2^ test.

Larval locomotion analysis was performed using a frustrated total internal reflection (FTIR) based tracking system (FIM, University of Münster)^63^. Five freely moving larvae per trial were placed on a 1 % agar plate (13x13 cm, or circular ø 10 cm arena) and video-captured with a CMOS camera (ac2040-25gm, Basler, Ahrensburg, Germany). During free locomotion, eACRs were activated by illumination with 525 nm light from a RGB-BL-S-Q-1R LED backlight (Phlox, Aix-en-Provence, France) for Aurora, or 460, 470 and 595 nm light from a pE-4000 (CoolLED) coupled to a light guide with custom collimator lenses for Phobos and Phobos^CA^. Animal locomotion was tracked with 10 frames/s for up to 210 s and then analyzed using FIMtracking software (FIM, University of Münster). Each genotype was tested multiple times on different days and data from all trials were combined. For analysis, only animals displaying continuous locomotion before the light stimulus were kept. Locomotion velocity was analyzed over time and was displayed with 1 s moving average. For comparison, velocities were averaged over a 5 s interval, each before, during and after light mediated activation of eACRs.

To compare the effect on body wall muscle contraction of inhibitory eACRs with excitatory ChR2^XXL^ ^41^ we analyzed the larval area change before and after light activation using FIMtracking software (FIM, University of Münster). Animal size was averaged over a 5 s interval before and after light activation for analysis. Statistical significance was calculated by ANOVA followed by a Sidak’s multiple comparisons test for repeated measurements or a (non-parametric) Kruskal-Wallis test followed by a Dunn’s multiple comparisons test for comparisons between groups.

Optogenetic activation/inactivation with Phobos^CA^ and CsChrimson was performed using the same setup as for larval locomotion. 10-20 freely moving larvae per trial were placed on a 1 % agar plate (circular ø 10 cm arena) and video-captured. During free locomotion, Phobos^CA^ expressed in A08n neurons (82E12-Gal4, UAS-PhobosCA) was activated by illumination with 460 nm light from a pE-4000 (CoolLED) light source. 30 s later, CsChrimson (27H06-Lex, LexAop-CsChrimson) was activated with a 635 nm light pulse of 5 s. Red light activation of CsChrimson without Phobos^CA^ activation or blue and subsequent red light illumination of animals expressing only CsChrimson served as controls. Videos were analyzed offline for animals showing nociceptive rolling behavior. Statistical significance was calculated by a chi^2^ test.

### Immunohistochemistry of larval brains and neuromuscular junctions (NMJs)

3rd instar larval brains were prepared as previously described^39^. For NMJ visualization, larval fillets were dissected in PBS and pinned down on Sylgard plates (Dow Corning, Midland, MI) with minutien pins (Fine Science Tools, Heidelberg, Germany). Animals were cut open dorsally and internal organs were removed while leaving the nervous system intact. Subsequently, larval brains or fillet preparations were fixed in 4 % formaldehyde/PBS for 20 min, blocked in PBS/0.3 % Triton X-100 containing 5 % normal donkey serum (Jackson ImmunoResearch Laboratories, West Grove, PA). Larvae expressing Aurora-Cerulean or Phobos^CA^-Cerulean in motor neurons (*ok371-Gal4*) were immune stained using a rabbit anti-GFP antibody (1:500, cat. no. A-11122, 1DB-ID: 1DB-001-0000868907, ThermoFisher Scientific). Secondary donkey anti-rabbit-DyLight488 and anti-HRP-Cy3 (NMJ surface visualization) antibodies were used at 1:300 dilution (cat. no.123-165-021, DB-ID:1DB-001-0000865678, Jackson ImmunoResearch). Larval brains and NMJs at muscle 6/7 were visualized by confocal microscopy with a 40x/NA 1.3 oil or 20x/NA 0.8 air objective, respectively (Zeiss LSM700, Carl Zeiss).

### Statistics

All statistical analyses were performed using GraphPad Prism 6.0 or Origin 10.5. Data were tested for normal distribution by D’Agostino & Pearson omnibus normality test. Normally distributed data were tested for significant differences (*p< 0.05, **p< 0.01, ***p< 0.001 and ****p< 0.0001) with one-way repeated-measures analysis of variance followed by Tukey’s, Dunnett’s or Sidak’s multiple comparisons test. Not normally distributed data were tested with the nonparametric Kruskal-Wallis test followed by Dunn’s multiple comparisons test. Data are presented as mean ± standard error of the mean (SEM). No statistical measures were used to estimate sample size since effect size was unknown. Given n numbers represent biological replicates (i.e. HEK-cells, neurons, *Drosophila* larvae). For nociceptive *Drosophila* experiments, but not for HEK-cell measurements and hippocampal neuronal recordings, investigators were blinded to the group allocation during the experiments. Data analysis was done by expert investigators who did not carry out the experiments. In addition, unsupervised analysis software was used if possible to preclude investigator biases. All experiments were done with interleaved controls and treatment groups were mixed, where possible.

### Data availability

The authors declare that all data and code supporting the findings of this study are included in the manuscript and its Supplementary Information or are available from the corresponding authors on request. The plasmids used in this study are deposited with the Addgene plasmid repository.

## Acknowledgements

We thank Edward Boyden (TcChR, TsChR, *Co*ChR, Chronos and Chrimson), Johannes Vierock (C1C2) and the late Roger Y. Tsien (ReaChR) for providing plasmids encoding for CCRs. We further thank Ivan Haralampiev, Thomas Korte and Andreas Hermann for help with confocal microscopy, Iris Ohmert and Sabine Graf for hippocampal slice cultures, Benjamin S. Krause for discussions and Kathrin Sauter, Maila Reh, Altina Klein and Tharsana Tharmalingam for technical assistance.

This work was funded by grants from the European Research Council (ERC-2016-StG 714762 to J.S.W), the German Research Foundation (SPP 1926 to P.S. & J.S.W.; FOR 2419 to J.S.W. and T.G.O., SFB1078 B2, FOR 1279, SPP 1665 to P.H. and T.G.O.). P.H. is Hertie Senior Professor for Neuroscience and supported by the Hertie Foundation.

## Author contributions

J.W., T.G.O., P.S., P.H. and J.S.W. conceived the study and planned experiments, J.W., S.R.R., J.T., F.T., C.G. and J.S.W. performed the experiments, J.W., S.R.R., J.T., F.T., C.G., P.S. and J.S.W. analyzed the data. J.W. and J.S.W. wrote the manuscript with contributions from all authors.

## Competing interests

The authors declare no competing financial interests.

